# Light induced changes in starry flounder (*Platichthys stellatus*) opsin expression and its influence on vision estimated from a camouflage-based behavioural assay

**DOI:** 10.1101/2020.07.30.228627

**Authors:** Tom Iwanicki, Cliff Haman, Amy Liu, John S. Taylor

## Abstract

Correlations between variation in opsin expression and variation in vision are often assumed but rarely tested. We exposed starry flounder (*Platichthys stellatus*) to either broad spectrum sunlight or green-filtered light in outdoor aquaria for seven weeks and then combined digital-PCR and camouflage experiments to test two hypotheses: i) short-wavelength sensitive opsin expression decreases in a green light environment, and ii) if observed, this change in opsin expression influences colour vision as estimated using a camouflage-based behavioural assay. Of the eight visual opsins measured, *Sws1* (UV sensitive) and *Sws2B* (blue sensitive) expression was significantly lower in fish exposed to green light. However, opsin expression in fish transferred to an arena illuminated with white LED light for three hours after the green light treatment did not differ from broad spectrum controls. Changes in opsin expression in response to artificial light environments have been reported before, but rapid changes over three hours rather than days or weeks is unprecedented. We did not observe a significant difference in a flounder’s camouflage response based on light environment, although broad spectrum fish increased and green-filter fish decreased the pattern contrast when on the blue-green substrate, and this difference approached significance. This pattern is intriguing considering green-filter fish expressed fewer UV and blue opsins and we recommend increased statistical power for future experiments. Together, our results show that starry flounder opsin expression changes rapidly in response to changes in light environment, however, there is no apparent effect on their visually mediated camouflage.

## Introduction

The ability to detect and discriminate among different wavelengths of light depends on the diversity of opsins in the photoreceptors of the retina. Humans are considered to be trichromatic, expressing short-wavelength sensitive (*OPNSW1*), middle-wavelength sensitive (*OPN1MW*), and long-wavelength sensitive (*OPN1LW*) opsin genes in retinal cone cells, and rhodopsin (*RHO*) in rod cells, which are used for scotopic (dim light) vision (1). Teleost fish typically have many more visual opsins than other vertebrates (2,3), largely as a result of lineage-specific tandem duplications (2,4). The advantage of large opsin repertoires, however, is not clear; humans can discriminate between colours (wavelengths) that differ by less than one nm over much of the visible spectrum (5).

It may be that a large visual opsin repertoire is a ‘toolkit’, with subsets of opsins used at different stages of development, in different light environments, or indeed in particular regions of the retina. Several authors have reported observations consistent with this hypothesis: developmental variation (6), geographic distribution based on light environment (7), and regions of the retina (8,9). Despite correlations between opsin repertoire or expression patterns and the light environment, clear connections between variation in opsin expression and visual performance have rarely been demonstrated outside of humans. Smith et al. (2012) manipulated opsin expression in Lake Malawi Cichlids and showed that LWS expression variation account for about 20% of the observed variation in optomotor response (10). In addition Sakai et al. (2016) found that *Lws-3* expression increased in guppies grown in orange light and that fish with higher levels of *Lws-3* expression had higher visual sensitivity to 600 nm light (11). Conversely, Wright et al. (2020) found colour perception plays an important role in female cichlid mate preference but opsin expression was only weakly correlated and a direct causal link between expression and behaviour was lacking (12).

Adaptive camouflage in Pleuronectiformes was first described by Sumner (1911): turbot (*Rhomboidichthys podas* and *Lophopsetta maculata*), summer flounder (*Paralichthys dentatus*), and winter flounder (*Pseudopleuronectes americanus*) changed patterns over a period of days in response to various mottled sandy substrates and checkerboards (13). Juvenile plaice (*Pleuronectes platessa*) change colour more quickly (14) and many bothids (e.g., left-eye flounder) can camouflage to environment cues in seconds (15). Rapid changes in camouflage are based on visual cues and can be used to infer visual performance. Flounder camouflage match natural substrates well when modeled to mono-, di-, and tri-chromatic visual systems (16), however, here we elected to use checkboards (as in Mäthger et al. 2006 (17)) to elicit an exaggerated pattern response. We held starry flounder (*Platichthys stellatus*), a flatfish possessing eight visual opsin genes (9), in outdoor aquaria exposed to either sunlight or green-filtered light anticipating opsin expression would adjust to each environment as in Fuller & Claricoates (2011) (18). We quantified the starry flounder’s camouflage response to measure the effect of opsin expression variation on vision.

## Materials and Methods

### Fish Collection and Light Exposures

Fish were collected by beach seine at low tide between 10:00 and 12:00 in May 2015 at Willows Beach, Victoria, British Columbia, Canada. The seine net was deployed from a small aluminum boat at a depth of approximately 3 m, or by hand at approximately 1 m depth. Sixteen starry flounder were transported to the Outdoor Aquatics Unit at the University of Victoria, and held under ambient light in 12°C recirculating seawater. These fish (TL = 172±28mm, m = 74.8±37.8g) were then distributed among eight experimental enclosures (Fig 1) on July 22, 2015. Fish were exposed to either green-filtered or broad spectrum light for seven weeks. Light transmission (%) for each filter (broad spectrum: Roscolux #3410; green: Roscolux #90) was measured using Ocean Optics QE Pro at -10°C (sensor), integration time 100 µs, average of 3 for each spectrum, boxcar width (2), and electric dark current correction. Over the course of seven weeks they were fed krill at 1% body weight per day, adjusted weekly at an estimated 2% specific growth rate. Feeding occurred once daily through a 2-cm hole in the lid that was otherwise sealed by a black rubber stopper to inhibit non-filtered light from entering the tanks. All procedures were approved by the University of Victoria Animal Care Committee, which abides by regulations set by the Canadian Council for Animal Care.

**Fig 1.**
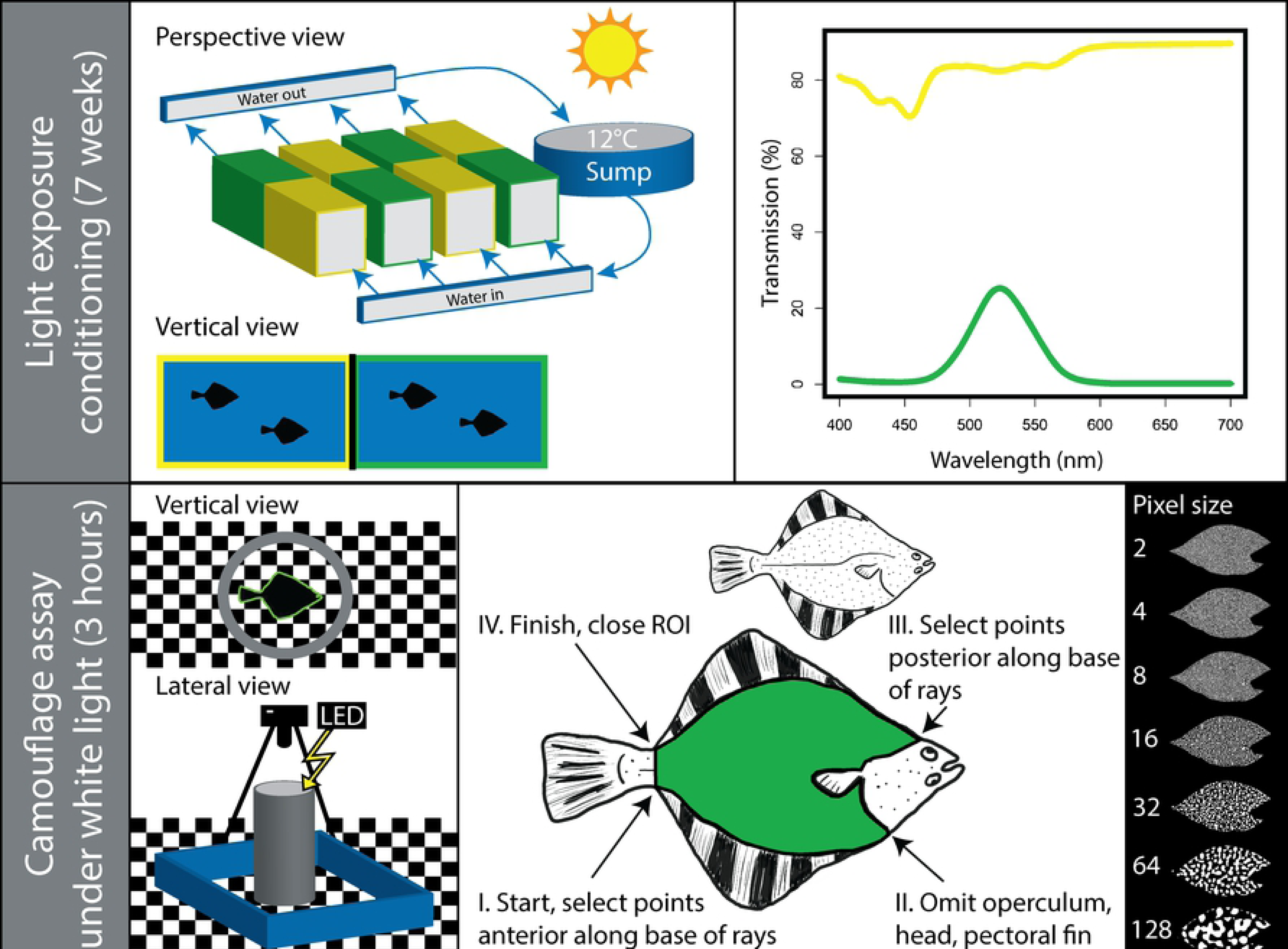
Schematic of the study design. Upper panels, Light exposure conditioning (7 weeks): Four tanks (*lwh* = 49”×19”×25”) were maintained outdoors on a 12°C closed, recirculating sea water system. Each tank was partitioned with an opaque plastic sheet with perforations to allow sea water to cycle through, but limit light transmission between treatments. Half of each tank was wrapped in Roscosun 1/8 CTO cinematic gel filter (Roscolux #3410) and the other half in Dark Yellow Green cinematic gel filter (Roscolux #90). Two fish were held in each enclosure, one fish was immediately euthanized after 7 weeks exposure, and the other fish proceeded to the camouflage assay for three hours prior to being euthanized. Light transmission (%) for each filter (top right: Roscolux #3410, yellow line; Roscolux #90, green line) was measured using Ocean Optics QE Pro. Bottom panels, Camouflage assay under white light (3 hours): one fish per enclosure was randomly selected for the behavioural assay. The individual was placed sequentially on five different substrates, illuminated with white LEDs, for 30 minutes per substrate. RAW images were captured on a Nikon DSLR and camouflage analysis was run on randomly selected images with methodologically selected regions of interest (ROI). Each ROI was “filtered” through seven bandpass filters (bottom right panel) and the pixel energy at each spatial scale is measured to quantify the camouflage pattern.

### Behavioural Assay

The behavioural assay was conducted in a dark room with a sea-tray table containing 12°C seawater on a recirculating system. The arena was comprised of a 30 cm diameter, 50 cm tall plastic tube with four XLamp Neutral White 4000K LEDs (Cree, Inc.) mounted on top of the tube in pairs at opposite ends. Neutral white foam core was used to reflect light into the arena to reduce hotspots and shadows. Laminated checkerboard substrates were inserted vertically and horizontally on the inside of the tube. The substrates were designed in Adobe Photoshop CS6. The colour space used was CMYK US Web Coated SWOP v2. The printer was an Epson Stylus Pro 9890 with Epson UltraChrome K3^®^ Ink package, and substrates were printed on Epson Premium Luster Photo Paper (206). A total of five substrates were printed, one uniform grey and four checkerboards (i.e., blue-green, blue-red, red-green, and black-white). The pigments used for the colourful checkerboards were selected with the saturation resulting in equal percent-reflectance of total photons. Equal reflectance of photons resulted in reflected light intensity being equal across the checkerboard, limiting the spectral signal from the checkerboards to hue (or wavelength), and reducing the role of contrast (intensity) as an explanation for a camouflage response. Percent-reflectance was calculated using an USB2000 Spectrophotometer (OceanOptics Inc.) and the reflectance software in OOOIBase. Images were captured using a Nikon D3100 Digital SLR camera and an AF Nikkor 50mm 1:1.4D lens. The camera settings remained constant for the duration of the experiment (aperture = F5.6, shutter speed = 1/5”, ISO = 200, exposure compensation = +0.7). The camera was mounted on a tripod approximately 1.5 meters above the behavioural arena.

Beginning on September 9, 2015, two fish from a randomly selected enclosure were selected on each of eight days. Diel rhythms in opsin expression have been previously reported in fish (19,20), therefore, individuals were chosen by balanced design, alternating between broad spectrum and green-filtered enclosure at the same time daily. One fish was selected (by coin toss) for the behavioural trial and transferred to the experimental arena. Fish were acclimated to the arena for 30 minutes before the five-substrate assay began. The other fish was euthanized and whole retinas dissected to provide baseline opsin expression, unaffected by the bright white lighting in the behavioural experiment. All behavioural trials began at 9:00AM to control for the effects of circadian rhythms. Fish were exposed to five substrates in the following order at intervals of 30 minutes (starting time in brackets): acclimatization period (9:00AM), grey (9:30AM), blue-green (10:00AM), blue-red (10:30AM), red-green (11:00AM), and black-white (11:30AM). Photos were captured at intervals of 30 seconds (60 photos per substrate, 300 photos total per individual).

### Image Analysis

Images were captured in RAW file format. A total of six randomly selected images were analysed per substrate per individual (6 images per substrate, 30 images per individual, 240 images total). Multispectral images were generated from RAW files and analyzed using the Image Calibration and Analysis Toolbox (21). All images were calibrated to a standard (PTFE sheet). Fish camouflage response was characterized from a cropped image. Cropping was performed by a person who was not aware of the study design. Image cropping followed a standard protocol, in short: a polygon representing the region of interest (ROI) was created starting at the base of the anal fin near the caudal peduncle. Points of the polygon were selected at the base of every third ray of the anal fin extending anterior the caudal fin. The pelvic fin, operculum, head, and pectoral fin were excluded from the polygon. The polygon extended posteriorly along the base of the dorsal fin (points at every third ray) back to the caudal peduncle and the polygon was closed off completing the ROI.

Granularity analysis similar to that used to quantify cuttlefish camouflage (22) and avian egg pattern (23) was used to get a single measurement for camouflage pattern; cropped images were filtered using each of seven spatial frequency bands, or bandpass filters (i.e., 2, 4, 8, 16, 32, 64, 128 pixels). The pattern of individual fish was estimated using the standard deviation of luminance, which measures the overall contrast within an image modelled to human vision. Higher standard deviation of luminance equates to more light-and-dark contrasting patterns (i.e., disruptive or mottle camouflage), whereas low values equate to low pattern contrast (i.e., uniform camouflage).

Two-way repeated measures ANOVA was run in R version 3.2.4 using the “nlme” package and Tukey multiple comparisons was run using the “multcomp” package. Analyses were based on standard deviation of luminance from a total of eight fish, held for seven weeks in either broad spectrum sunlight or green-filtered light, on five chromatically different substrates. The mixed effects model tested was: camouflage ~ light environment + substrate + light environment × substrate + (1|individual) + ε.

### RNA isolation and digital PCR

Eyes were removed and a razor blade was used to cut the cornea exposing the lens and retina. The lens was removed and the retina extracted. Retinas were frozen in liquid Nitrogen and stored at -80°C. Retinas were then homogenized in TriZol (Invitrogen) with zirconia beads using a mini beadbeater (BioSpec products) for 30 seconds. RNA was isolated following the TriZol manufacturer’s protocol, with slight modification. The RNA pellet was washed twice (rather than once) with >75% ethanol. DNA, if present, was digested using RNase-free DNase I (ThemoFisher Scientific, EN0521). Total RNA was quantified using Qubit^®^ RNA Broad Range Assay Kit (ThemoFisher Scientific, Q10210). 1 µg of RNA from each sample was reverse-transcribed in 40 µl using iScript™ cDNA Synthesis Kit (BioRad).

Digital-PCR (dPCR) was run on QuantStudio^®^ 3D Digital PCR System (Life Technologies) using locus-specific primers and TaqMan probes for all eight visual opsins found in the starry flounder transcriptome (Table 1). Opsins were multiplexed using FAM and VIC reporter dyes. cDNA, primers, probes, and master mix were loaded onto a QuantStudio^®^ 3D Digital PCR 20K v2 Chip and sealed with immersion oil to prevent evaporation. After equilibrating at room temperature for 15 minutes, PCR was performed on a ProFlex™ 2x Flat PCR System (step 1: 94°C × 30 sec; step 2: 55°C × 2 min, 94°C × 30 sec (39 cycles); step 3: 55°C × 2 min, 10°C hold). Chips were read using the QuantStudio^®^ 3D Digital PCR instrument. Sample concentrations were adjusted to ensure that transcripts per microliter fall within the digital range of the 3D system (i.e., 200 – 2000 copies•µl-1). cDNA template varied from 0.1 to 100 ng per chip. Opsin expression was normalized using the alpha subunit of transducin (*Gnat2*), the G-protein activated by cone opsins (19). Patterns in expression were tested using a paired student’s t-test. All statistical tests were evaluated at α = 0.05 level of significance.

**Table 1:**
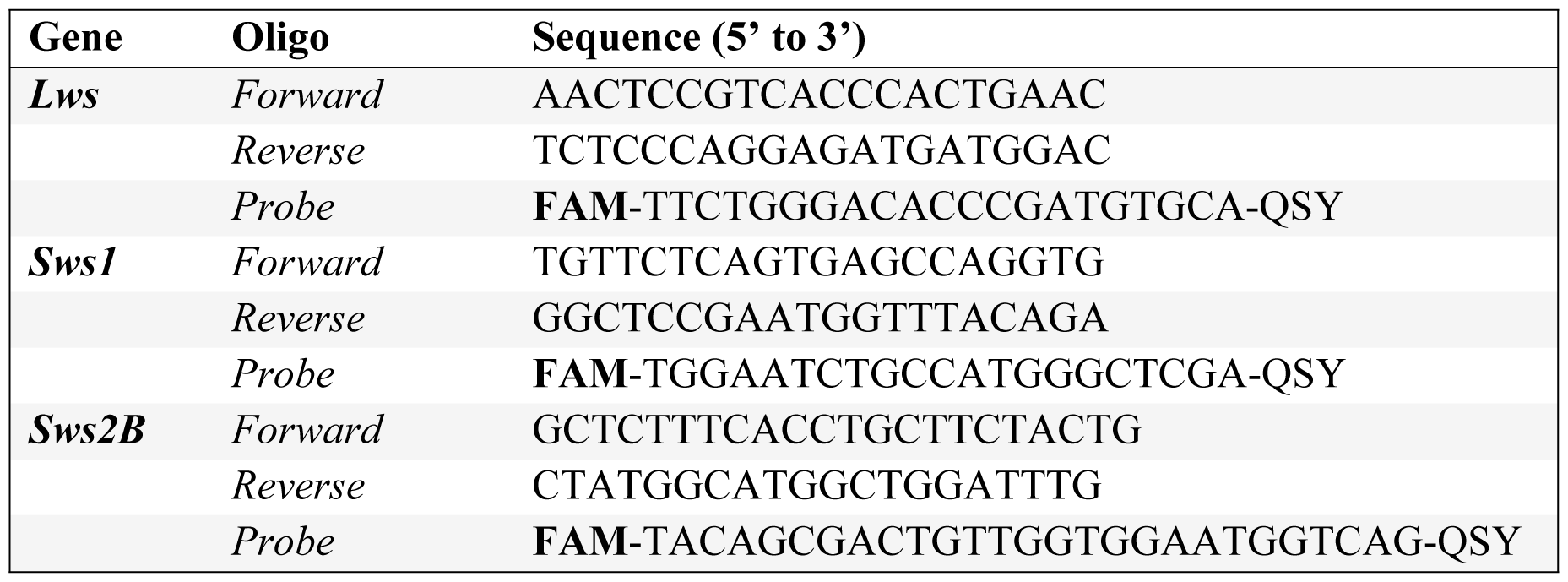
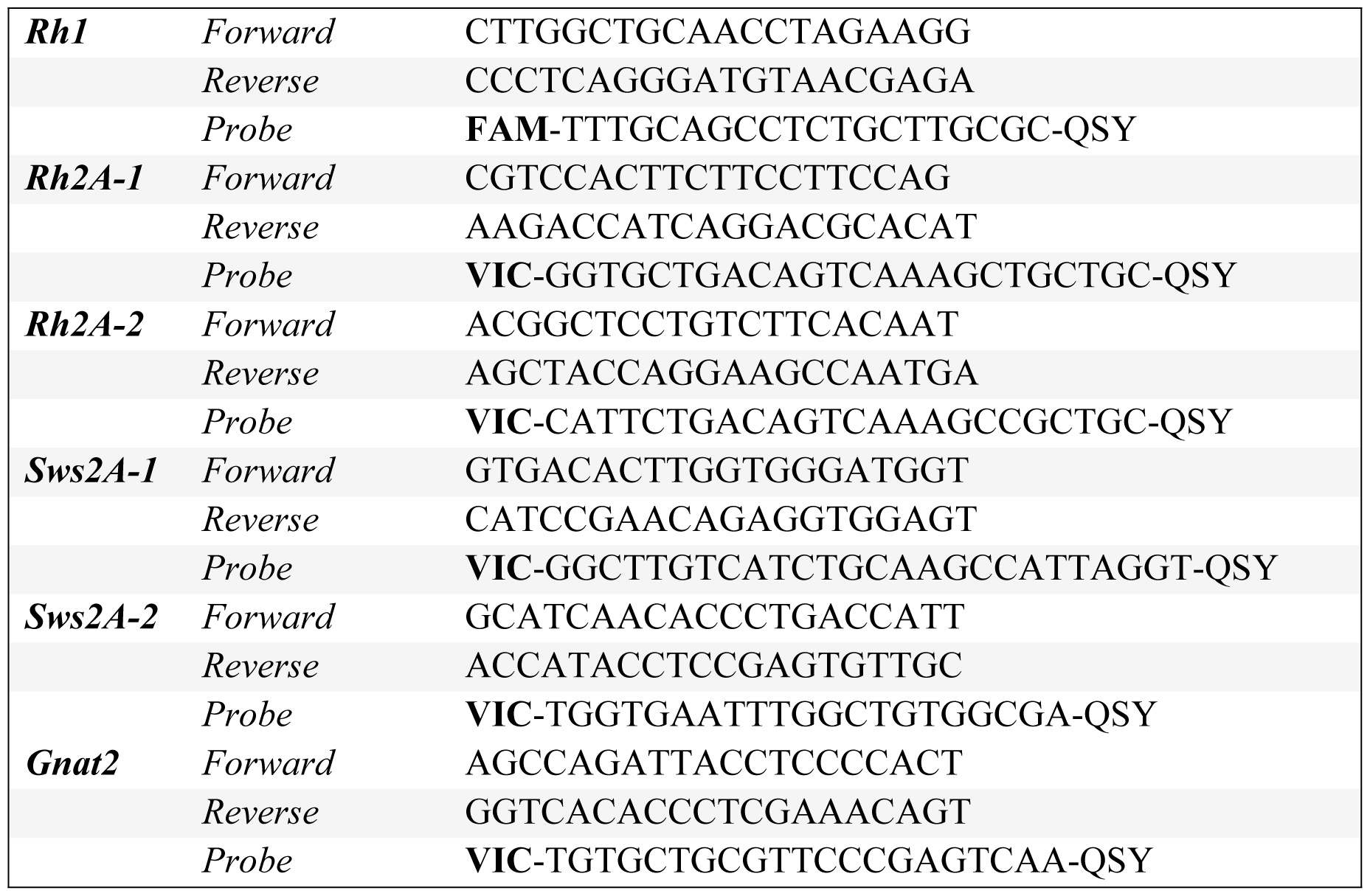
Primers and TaqMan probes used for starry flounder digital-PCR.

## Results

### Experimental animals

Fish varied in size but this variation was distributed among treatment and control aquaria. Light treatment did not influence growth over the seven-week exposure (broad spectrum: ΔTL = 7.5±5.9 mm; green-filtered: ΔTL = 4±7.2 mm) or mass (broad spectrum: Δmass = 1.2±9.3 g; green-filtered: Δmass = 0.9±9.2 g). There was no statistical difference in length and mass of baseline fish (i.e., those immediately euthanized after seven weeks of conditioning) and fish used in the behavioural assay (i.e., fish exposed to bright white LED for 3 hours after conditioning) (baseline: TL = 165.20±24.20 mm and mass = 61.60±29 g; time 3 hours: TL = 190.80±29.42 mm and mass = 89.97±36.20 g; TL: t = -1.8925, p = 0.08006 and mass: t = -1.7296, p = 0.1067).

### Image analysis

Fish patterns changed in response to the substrate. The mixed effects model for the camouflage indicated substrate (checkerboard) was significantly associated with the camouflage pattern (F = 4.552, p = 0.0071). When placed on a blue-green substrate, fish exposed to broad spectrum light displayed greater pattern contrast than fish from the green light treatment, but the difference was not significant (F = 5.767, p = 0.0532) and the interaction between substrate and light environment did not significantly influence camouflage (F = 1.209, p = 0.3327). Tukey multiple comparisons indicated that the pattern of fish from broad spectrum light on the black-white substrate was significantly different than on: i) broad spectrum, blue-red substrate (z = 3.135, p = 0.0493), ii) green-filtered, grey (z = 3.506, p = 0.015), iii) green-filtered, blue-green (z = 3.876, p < 0.01), and iv) green-filtered, blue-red (z = 3.338, p = 0.0264). Green-filtered, black-white was significantly different than green-filtered, blue-green (z = 3.3132, p = 0.0498). Overall, contrast (i.e., black-white substrate) results in the greatest pattern change in both treatments and the effect of light environment approached significance, based on Tukey multiple comparisons the difference was driven by differential camouflage response on the blue-green substrate (z = - 3.110, p = 0.0536) (Fig 2).

**Fig 2.**
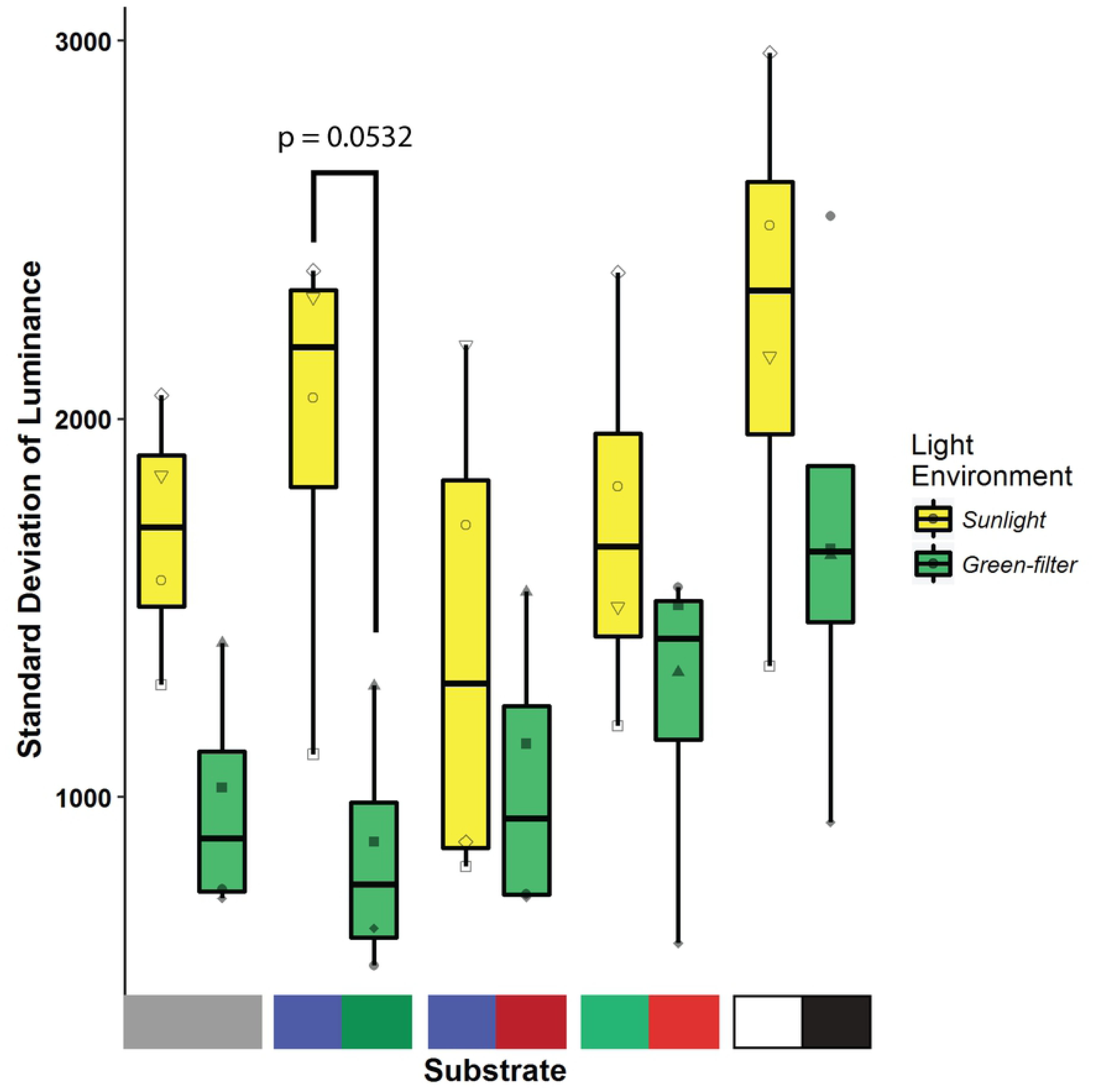
Camouflage pattern measured as the standard deviation of luminance across seven spatial frequency bands (granularity analysis) of starry flounder (shapes represent individuals) on five different substrates as depicted on the x-axis (left to right: grey, blue-green, blue-red, green-red, and black-white). Fish were conditioned for seven weeks to either broad spectrum sunlight (yellow bars) or green-filtered light (green bars).

### Digital-PCR

Fish that were immediately euthanized after being removed from the 7 week light treatment (e.g., the ‘baseline fish’) had significantly different opsin expression; individuals held in green-filtered light had lower expression of UV sensitive (*Sws1*) and short-wavelength sensitive (*Sws2B*) opsins compared to those exposed to broad spectrum light (student’s t-test, t = 3.9414, p = 0.01121 and t = 1.1458, p = 0.004792, respectively) (Fig 3). Opsin gene expression levels were the same in fish from the broad spectrum and green-filtered light exposure that were transferred to the behavioural arena and exposed to white LED light for three hours (i.e., the duration of the behavioural assay) (Fig 3).

**Fig 3.**
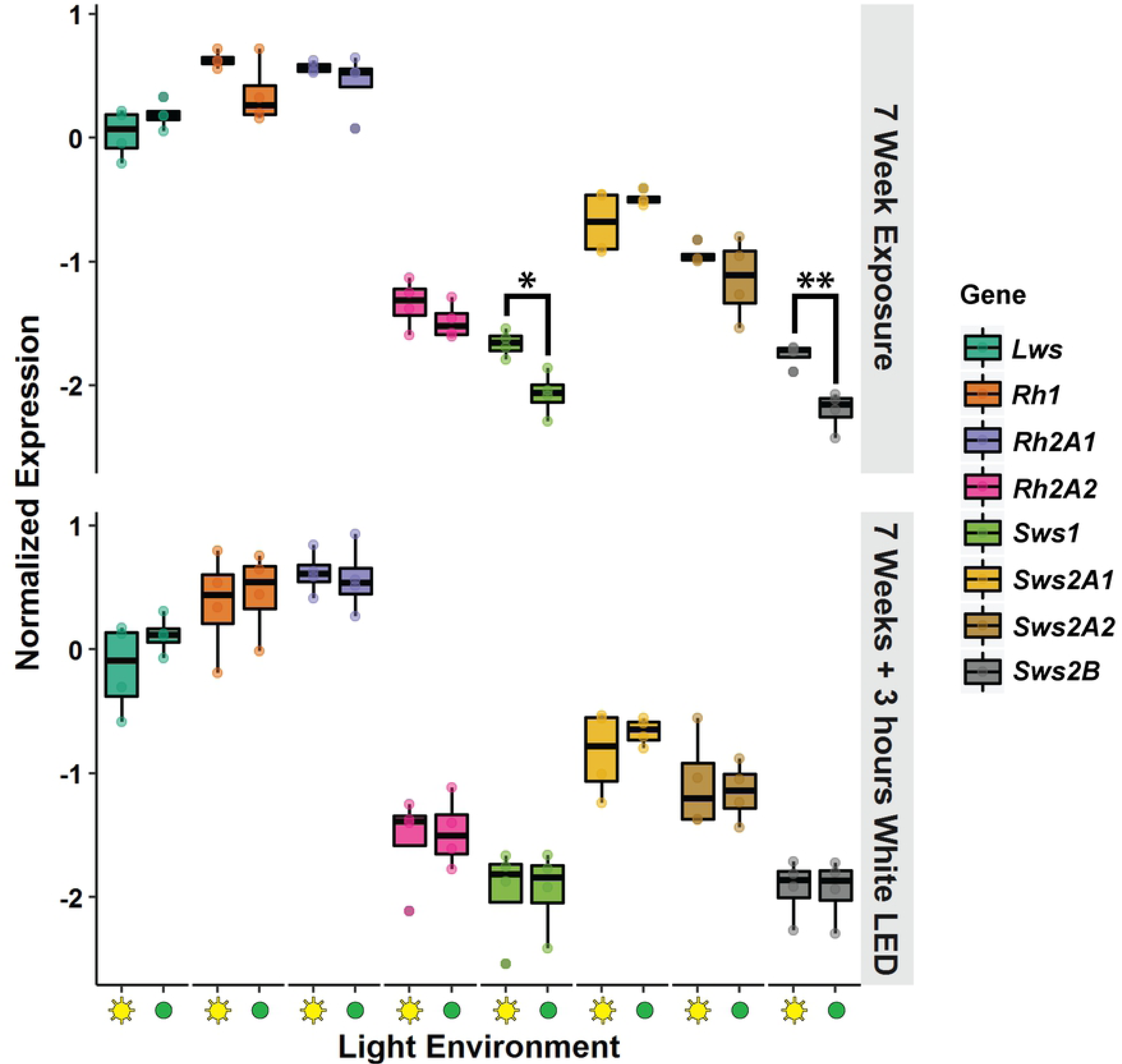
*Gnat2* normalized opsin expression of starry flounder held in either broad spectrum sunlight (x-axis, C) or green-filtered sunlight (x-axis, G) for seven weeks. Fish (n=4) were euthanized immediately after being removed from the light environments (the “7 Week Exposure” panel, top) or 3 hours after being transferred to the behavioural arena (the “7 Weeks + 3 hours White LED” panel, bottom) illuminated with four white LED lights (n=4). Asterisk denote significant differences between light exposure (*: p = 0.01121; **: p = 0.004792).

## Discussion

### Opsin expression plasticity in response to light environment

Transcripts of eight distinct visual opsins are expressed in the eyes of juvenile starry flounder. Microspectrophotometry data indicate that all are translated and that just one type of chromophore is used (9). We predicted opsin expression would be modified by a seven-week exposure to distinct light environments and that changes in opsin expression over that length of time would influence vision. We used a camouflage-based assay to assess visual performance. Opsin expression in the starry flounder retina did change in response to the light treatment, and then changed again within three hours under white LED light.

Experiments designed to influence opsin expression have succeeded in the past, but the time scale observed here is unprecedented. Killifish reared in clear or tea stained water were monitored over four weeks and opsin expression differences were observed within 1-3 days (18). Opsin expression from clear and tea stained water were concordant with natural killifish populations and suggest opsin plasticity is used to tune vision to the light environment (18,24). Here we show that *Sws1* (UV) and *Sws2B* (blue) expression was lower in the individuals that spent 7 weeks in an environment lacking wavelengths below 450 nm compared to those in broad spectrum light. The six other visual pigments have wavelengths of maximum absorbance within the light available in the green-filtered tank, and were expressed at the same level in all fish despite the overall light intensity being markedly different between the green and broad spectrum tanks. The difference in *Sws1* and *Sws2B* expression, induced by the absence of short-wavelength light, was lost after only three hours of exposure to white light in the camouflage trials. Although the white LEDs do not emit UV light, they do emit near-UV and blue light, which was enough to induce higher expression of *Sws1* and *Sws2B* opsins. Development can play a role in UV opsin expression. In Salmonids UV opsin is one of the first opsins expressed in the larval fish and is subsequently lost as they develop and transition into an active lifestyle following smoltification (25). However, the differences observed here were not due to ontogeny, as *Sws1* and *Sws2B* expression was not correlated with fish length or mass. Further, opsin expression changes occur more rapidly in single cones than double cones (26), and both opsins rapidly affected here are found in single cones.

The rapid plasticity of opsins on the order of hours, rather than days, has implications for the visual ecology of starry flounder and the study of opsin expression in natural populations more broadly. The changes could function as a way of tuning the retina to varying light conditions, rapidly setting the machinery in place to restructure the retina if novel light conditions persist. Increasing populations of photoreceptors sensitive to the light in the environment could improve visual sensitivity and confer benefits for predator avoidance and prey capture. Future studies should investigate whether plasticity is a persistent phenomenon throughout starry flounder ontogeny, or if the retina is plastic at certain stages of development. Juvenile starry flounder are found in shallow, nearshore waters and as adults descend to depths of more than 200 meters, but occasionally migrate kilometers up river (27). These three environments (e.g., coastal shallows, benthic depths, and river) are spectrally dramatically different, and a tunable retina even at later ontogenetic stages could be adaptive. Given the logistical constraints of sampling wild populations of fish, the rapid change in opsin expression has implications for studies moving forward. One must consider both the time until preservation and the light conditions one is sampling in. We recommend as standard practise to limit ambient light while collecting samples in the field and to perform dissections under red light. Where ever possible, gear that limits introducing novel ambient light should be used (e.g., a closing cod-end on a trawl net).

Varying light environments, driven by water depth or season, affect opsin expression in several species of damselfish, whereas other species appear to have more stable expression patterns (28). In stickleback, opsin expression is shifted toward longer wavelengths in freshwater populations relative to marine populations, and these shifts correlate with differences in the light available (29). Furthermore, there is evidence for local adaption to light among benthic and limnetic ecotypes within a lake. These aforementioned differences in opsin expression were maintained in laboratory rearing experiments under fluorescent light illumination, ergo stickleback opsin expression is primarily under genetic control, a result of standing genetic variation (29). Why some species appear to possess plastic opsin expression while others do not warrants further investigation.

A change in opsin expression may not immediately reflect the opsin proteins present in the outer segment of a cell. Photoreceptors are terminally differentiated; they are long lived, and the cellular components must be regularly turned over to prevent a loss of function (30). Outer segment membranes, the light sensitive region of a photoreceptor, are shed distally and the addition of new membranes at the base renews the outer segment components. In mouse, rat, and frog, radioactively labelled amino acids accumulated at the base of the outer segment within 24 hours. Furthermore, in rods the labelled amino acids proceeded as a “reaction band” to the distal point of the outer segment in approximately ten days (31). Similar observations were observed in rhesus monkey and cat cone cells (32). For the aforementioned reasons we made the duration of the initial light treatment (i.e., seven weeks) sufficiently long to allow for protein-level changes throughout the outer segments of the starry flounder retina. Additionally, we do not expect the three hour period in which opsin expression returned to baseline to be enough time to make functional changes at the protein-level. Protein-level changes are observed over longer periods of time, such as ontogenetic shifts in opsin expression in coho salmon resulted in protein-level shifts as measured by microspectrophotometry (34). Recently, we observed parallel opsin switches within the outer segments of double and single cones of starry flounder. The proportion of outer segments containing co-expressed opsins were greater in juvenile fish, with shorter wavelength-sensitive opsins expressed at the distal tip of the outer segment (33). Given the delay from expression of opsin mRNA to translated and localized visual pigments, the starry flounder used in the behavioural assay are better represented by the baseline opsin expression data than the data collected from their retinas after three hours under a white LED light. If the *Sws1* and *Sws2B* expression patterns were consistent with the baseline over seven weeks, then we predict that the opsin protein populations in the retina would reflect those changes. However, it would be valuable to complement future behavioural assays with immunohistochemistry, or a survey of the retina using microspectrophotometry to confirm our prediction.

### Empirical evidence for active camouflage in starry flounder

The behavioural assay presented here contributes to a large body of empirical evidence that flatfish can change their skin pattern in response to substrate changes. The fish tested showed noticeable changes within 10 seconds, and the body pattern was stable within minutes, indicating direct neural input. Camouflage in fish is the aggregate response of millions of chromatophores in the skin. Unlike cephalopod molluscs (35,36), fish camouflage involves the physical movement of pigment, rather than muscular contraction and expansion of the cell itself (37). Therefore, it is not surprising that starry flounder camouflage is relatively slow compared to the remarkably fast change observed in cephalopods and other more specialized flatfish, which in some cases can occur in as little as two seconds (15,38).

Light environment may have affected visual performance of starry flounder camouflaging on the blue-green checkerboard. Broad spectrum fish increased pattern contrast and green-filter fish decreased pattern contrast and the difference approached significance (Fig 2). Higher standard deviation of luminance equates to more light-and-dark contrasting patterns (i.e., disruptive or mottle camouflage), whereas low values equate to low pattern contrast (i.e., uniform camouflage). A fish camouflaging with greater contrast would indicate that the fish can see a difference between the blue and green checkers. The broad spectrum fish deployed more mottle camouflage compared to green-filtered fish, and may therefore detect a greater difference between the blue and green checkers. Differential visual performance on the blue and green substrate is supported by gene expression, with broad spectrum fish expressing more UV- and blue-sensitive opsins, possibly conferring greater visual sensitivity to short-wavelength light. These data suggest a positive correlation with the *Sws1* and *Sws2B* opsin expression and an ability to detect a difference between blue and green hues, however the behavioural experiment did not have power to detect a significant difference in camouflage. Power analysis indicates if light environment truly impacts performance, a sample size of 20 fish would be required to detect it reliably.

An alternative explanation is that the *Sws1* and *Sws2B* opsins are not main drivers of the camouflage response. As with double cones that contribute chiefly to luminance vision, motion detection (39), and polarization vision (40), specialization in either double or single cones may exist that contribute to camouflage not affected by variation in UV and blue opsin expression. Future studies using different filters attempting to change opsin expression in different photoreceptors (e.g, green and red cones) would be informative. Previous research on starry flounder found a preponderance of unequal double cones in the dorsal retina, which receives reflected light from the substrate (9), and it may be functionally important for the camouflage response. In cichlids, the pattern of expression in double cones was reversed dorso-ventrally in response to red light illumination from below (41). If similar bottom-up illumination of starry flounder “flips” the double cone opsin expression dorso-ventrally, we could test whether the unequal double cones play an important role in camouflage.

Camouflage may be visually mediated through achromatic channels (e.g., cuttlefish have only one visual pigment). The most pronounced pattern changes were observed on the black and white checkerboard. That is not to say colour vision is unimportant to camouflage behaviour. Gulf flounder (*Paralichthys albiguttata*) and ocellated flounder (*Ancylopsetta ommata*) of family Paralichthydae preferred to settle on blue and green substrates after being adapted to the same colour (42). Mäthger et al. (2006) created a series of checkerboards that were white and green, with the white checker position getting progressively darker until the final substrate was black and green (17). At some point the green checker matched the luminance of the grey checker, and cuttlefish deployed a uniform camouflage pattern when placed on top. Thus, cuttlefish camouflage is achromatic. A similar experimental design, or perhaps substrates designed based off of Ishihara plates, would be useful to confirm whether starry flounder camouflage is driven by luminance, colour, or both.

The experiment did not control for the difference in overall light intensity between the two environments. The green-filtered environment allowed approximately 12% sunlight through compared to 88% in the broad spectrum environment. Although it is possible the differences in opsin expression could be due to light intensity, the evidence contrary to that point is two-fold. One, the opsins that were expressed significantly lower correspond to the wavelengths of light omitted by the filters. Two, the other opsins were not noticeably affected by the significantly lower light intensities in the green-filtered tank. Our rational behind selecting the green filter was that it approximated the light environments starry flounder encounter at depths in the turbid, coastal waters around Vancouver Island, Canada. With that said, follow-up studies that match light intensity, but vary colour, would enhance the present study.

### Concluding remarks

We found significantly greater UV and blue opsin expression after seven weeks in starry flounder held under broad spectrum light compared to green-filtered light. Surprisingly, that difference was lost after three hours under white LED light, indicating more rapid plasticity in opsin expression than previously reported. The timescale of change has relevance to both the visual ecology of fishes and the logistics and possible bias of studying opsin expression in natural systems. By using starry flounder’s visually-mediated adaptive camouflage, we were able to quantify visual performance on a variety of substrates, though we did not find statistically significant differences among fish from different light environments and recommend greater statistical power for future behavioural experiments.

## Acknowledgements

We would like to thank Chris Secord for donating materials and Mike Delsey for assistance constructing the behavioural arena, Ronnie Duke and Norm Johnson from the UVic Aquatics Facility for helping construct and maintain aquaria including husbandry, Will Duguid for help with field site reconnaissance and sample collections, and finally Dr. Geir Johnsen for assistance with optics.

